# Normal Aging in Mice is Associated with a Global Reduction in Cortical Spectral Power and Network-Specific Declines in Functional Connectivity

**DOI:** 10.1101/2021.06.09.447772

**Authors:** Asher J. Albertson, Eric C. Landsness, Michelle J. Tang, Ping Yan, Hanyang Miao, Zachary P. Rosenthal, Byungchan Kim, Joseph C. Culver, Adam Q Bauer, Jin-Moo Lee

## Abstract

Normal aging is associated with a variety of neurologic changes including declines in cognition, memory, and motor activity. These declines correlate with neuronal changes in synaptic structure and function. Degradation of brain network activity and connectivity represents a likely mediator of age-related functional deterioration resulting from these neuronal changes. Human studies have demonstrated both general decreases in spontaneous cortical activity and disruption of cortical networks with aging. Current techniques used to study cerebral network activity are hampered either by limited spatial resolution (e.g. electroencephalography, EEG) or limited temporal resolution (e.g., functional magnetic resonance imaging, fMRI). Here we utilize mesoscale imaging of neuronal activity in *Thy1-*GCaMP6f mice to characterize neuronal network changes in aging with high spatial resolution across a wide frequency range. We show that while evoked activity is unchanged with aging, spontaneous neuronal activity decreases across a wide frequency range (0.01-4Hz) involving all regions of the cortex. In contrast to this global reduction in cortical power, we found that aging is associated with functional connectivity (FC) deterioration of select networks including somatomotor, cingulate, and retrosplenial nodes. These changes are corroborated by reductions in homotopic FC and node degree within somatomotor and visual cortices. Finally, we found that whole cortex delta power and delta band node degree correlate with exploratory activity in young but not aged animals. Together these data suggest that aging is associated with global declines in spontaneous cortical activity and focal deterioration of network connectivity, and that these reductions may be associated with age-related behavioral declines.

## Introduction

Normal aging is associated with declines in cognition across many domains including executive function, memory, and language production(1-3). Aging is also associated with reductions in sensorimotor function including slower performance of visuomotor tasks(4), reduced walk speed(5), declines in motor planning and dexterity(6), and worsened coordinated movement(7). These functional declines happen in concert with significant cerebral changes. Brain weight(8), myelinated fibers(9), and cortical thickness(10) all decline in relatively linear fashion during normal aging. Interestingly, these changes are not correlated with significant neuronal loss, but are rather associated with synaptic changes(11). Multiple studies in primate and rodent models have demonstrated age-related changes in dendritic length(12), spine number and type(12), synaptic dynamics(13), as well as short(14) and long term synaptic plasticity(15). White matter tracts also decline with age. Fiber tractography reveals that white matter tract integrity peaks in midlife before gradually diminishing with age(16-18).

The mechanisms by which the observed synaptic, neuronal, and white matter tract changes seen with aging lead to functional declines in behavior are poorly understood. One likely consequence of age-associated synaptic changes is disruption of neuronal network activity. Deterioration of large-scale cortical networks with aging may bridge the cellular and molecular changes of aging with the behavioral decline observed in aging. Network level synchronous brain activity is hypothesized to coordinate information transfer and contribute to sensory and cognitive functions(19, 20). Previous human studies have shown that aging is associated with significant changes to brain spontaneous activity and these changes may be indicators of cognitive performance(21). Perhaps most compelling, normal aging is associated with a significant and linear decline in resting state spontaneous cerebral activity in slower frequency ranges (0.5-6.5Hz) and slow wave power in older individuals is a significant predictor of cognitive performance(22).

Functional connectivity across brain networks may offer another important link between age-related synaptic changes and behavioral declines. fMRI-based resting state functional connectivity (FC) studies have demonstrated significant effects of aging on network connectivity (reviewed by Damoiseaux JS(23)). Among other changes, aging is significantly associated with decreased connectivity in both the anterior and posterior components of the default mode network(24). Grady et al. further demonstrated reduced connectivity in the default mode network as well as increased inter-network connections for the frontoparietal control and dorsal attention networks in aged individuals during tasks(25). These and other studies strongly suggest that normal aging is associated with regional reorganization of network functional connectivity.

Prior studies of cortical network activity and connectivity have typically utilized either EEG or fMRI. While EEG studies have relatively high temporal resolution, they have significantly limited spatial resolution(26). The EEG signal is limited by nature in recording brain-derived field potentials through scalp and skull tissue which acts to smooth the recording and limit spatial resolution(27). This limitation is further exacerbated by variability in head size, skull thickness, tissue type, and body temperature(28). Conversely, fMRI offers better spatial resolution, but has limited temporal range with observations restricted to activity in the infraslow spectra (< 0.08 Hz)(29). Furthermore, the BOLD signal measured in fMRI studies represents a vascular, indirect measure of neuronal activity(30). This is an important confound because numerous studies have demonstrated age-related deteriorations in neurovascular coupling(31, 32).

Here, we seek to address these limitations by characterizing age-related changes in the cortical network activity and connectivity by performing direct mesoscale (whole cortex) imaging of neuronal network activity in young and aged *Thy1*-GCaMP6f transgenic mice. Projection neurons within these mice produce a robust fluorescent signal with fast activation and deactivation and high signal to noise ratio in association with suprathreshold neuronal activity(33-35). Use of this technique allows direct characterization of age-related neuronal network changes with high temporal and spatial resolution(36, 37). We chose to examine cortical networks via several measures including evoked and spontaneous activity. We examined network connectivity using seed-based and homotopic correlations as well as whole-cortex node degree. While we first observed a global decline in cortical spectral power, our observations of functional connectivity revealed a significant visual and somatomotor component, so we quantified spontaneous behavior in aged and young animals and correlated these levels to whole-brain spontaneous activity and network connectivity.

## Results

### GCaMP neuron density and somatosensory-evoked activation are similar between young and aged mice

We compared cellular and physiologic characteristics of *Thy1*-GCaMP6f mice at two different ages: Young mice (2 to 3 months) and aged mice (17 to 18 months). To ensure that differences between ages were not due to changes in the density of GCaMP-expressing neurons, we first quantified GCaMP expressing cells within the cortex of young (n=3) and aged animals (n=3) in primary motor cortex (M1) and barrel cortex (S1Bf) (Figure 1a & Figure 1b). We found no age-dependent differences in GCaMP positive cell density in layer 2/3 barrel (Young= 55.86 ± 4.36 cells/mm^3^; Aged= 54.12 ± 4.86 cells/mm^3^) and primary motor cortex (Young= 49.75 ± 1.51 cells/mm^3^; Aged= 55.86 ± 8.32 cells/mm^3^) or in layer 5 in barrel (Young= 193.77 ± 11.41 cells/mm^3^; Aged= 173.70 ± 6.1 cells/mm^3^) and primary motor cortex (Young= 160.61 ± 3.15 cells/mm^3^; Aged= 147.51 ± 10.72 cells/mm^3^) (Figure 1c).

**Figure 1.**
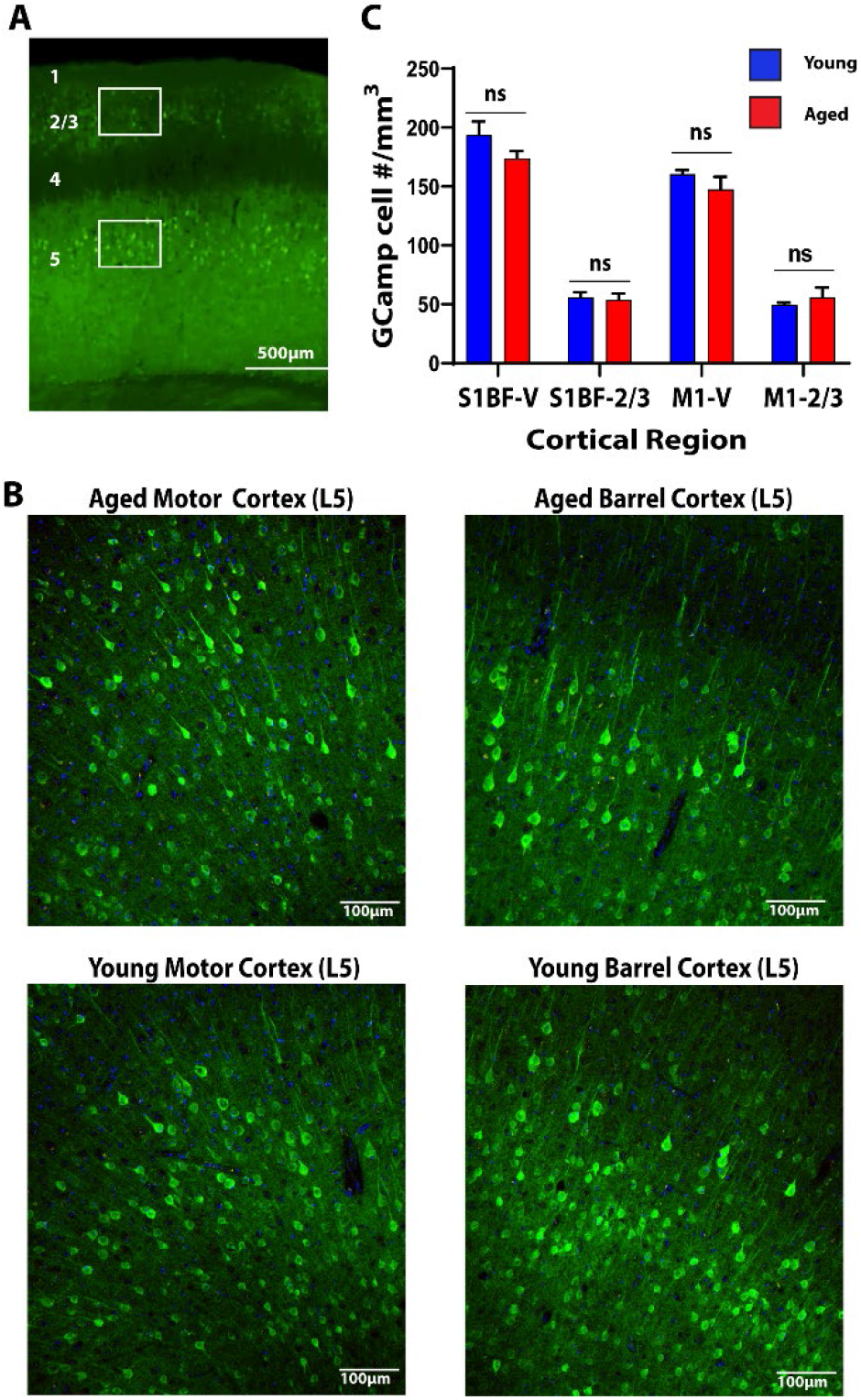
Young and Aged Mice have Similar Cortical GCaMP Expression. **A**. Low magnification view of the cortex. Numbers represent cortical layers. Boxes represent regions of interest in which counts were undertaken. **B**. Representative expression patterns from layer 5 motor and barrel cortex in aged and young animals. **C**. Quantification of GCaMP positive cell count in Layer 2/3 and Layer 5 barrel and motor cortices in aged (red) vs young (blue) animals.

To determine if differences between ages were due to changes in GCaMP activation, we next examined sensory-evoked activity. Somatosensory cortex activity was evoked via mild electrical stimulation of the right forepaw (Figure 2a). Stimulation evoked robust local somatosensory cortex activation in both young and aged mice (Figure 2b). Stimulation intensity was similar to prior experiments which demonstrated consistent, reproducible forepaw somatosensory cortex activation(38). We found no significant difference in either intensity (%ΔF/F) (Young= 0.23± 0.081; Aged= 0.27 ± 0.076) or area of activation (measured as pixels above threshold) (Young= 2143.7 ± 17.22; Aged= 153.4 ± 22.63) in aged vs. young mice (Figure 2c). Averaged traces of GCaMP signal intensity in the aged vs young cortex are shown in (Figure 2d). These data indicate that neither the density of GCaMP expressing neurons nor evoked GCaMP activity changed with aging.

**Figure 2.**
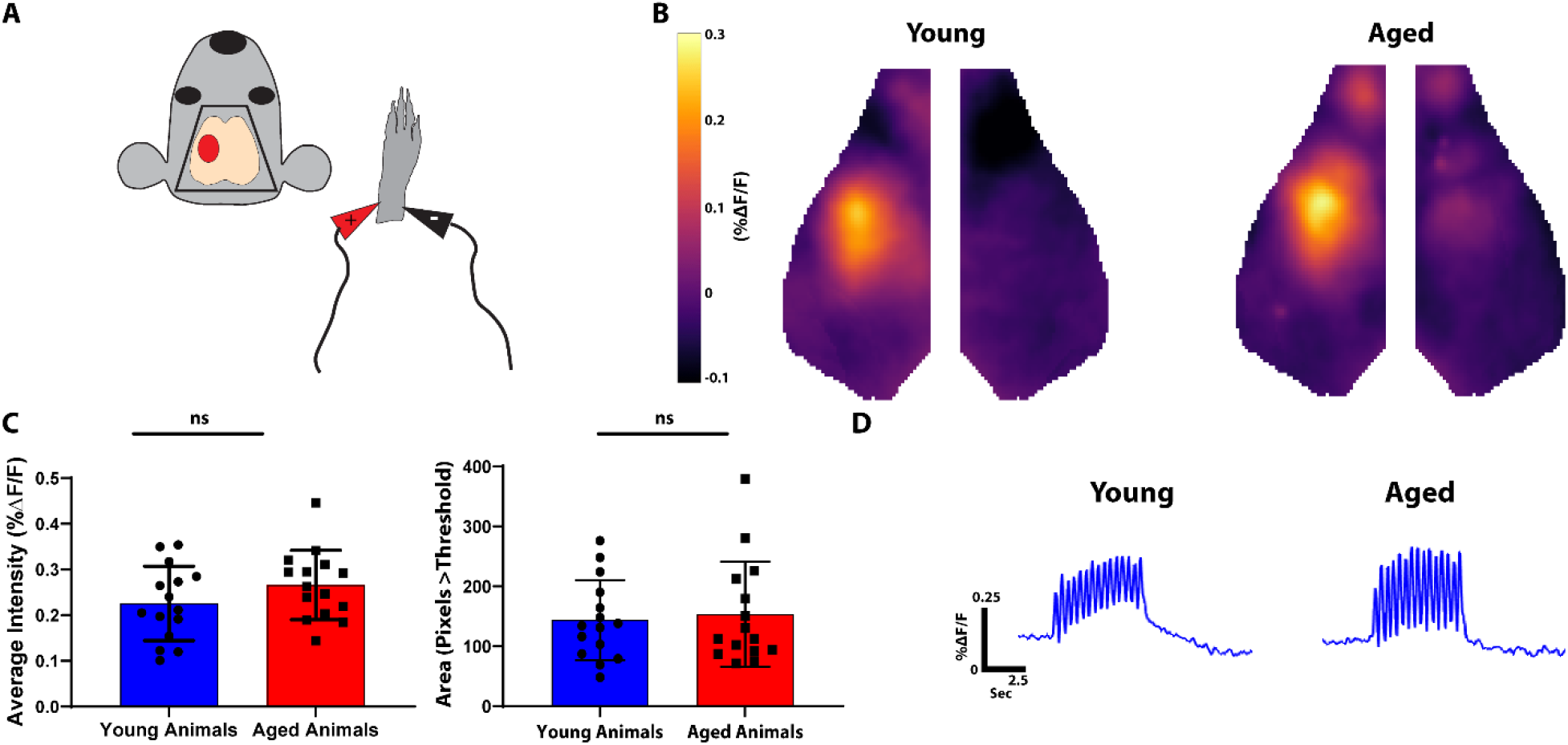
Young and Aged Mice have Similar Patterns of Evoked Somatosensory Cortex Activity. **A**. Diagram of experimental paradigm. The right forepaw was stimulated with mild electrical pulsations and contralateral somatosensory cortex activity was imaged (red oval). **B**. Average somatosensory activation in young (left) and aged (right) animals. **C**. Quantification of evoked somatosensory activity. The left graph represents average intensity within a somatosensory ROI. The right graph represents total area of activation as measured by pixels greater than threshold. **D**. Averaged fluorescence amplitude curves over time within a somatosensory ROI in young (left) vs. aged (right) animals.

### Cortical resting state power is globally reduced across all frequencies with aging

Compared to neural activity, hemodynamic measures of brain activity (e.g. fMRI, OISI) are relatively slow (the hemodynamic response function primarily acts as a low pass filter). Further, a variety of non-neurologic factors, including changes in neurovascular coupling and changes in vasoreactivity (Reviewed by Lu et al(39)), limit interpreting how local changes in neuronal activity relate to subsequent hemodynamics. We therefore sought to characterize spontaneous neuronal network activity using the high temporal and spatial resolution provided by GCaMP imaging. We examined band-limited power over the entire cortex across a wide range of frequencies (0.01 Hz-4 Hz). Global spontaneous neural fluctuations in aged mice were generally lower across the entire frequency range (Figure 3a). To quantify frequency-specific changes in neural activity, we subdivided activity into bins of equal relative bandwidth (2 octaves/bin) (Figure 3b). Aging was associated with a significant reduction in spontaneous neuronal activity across each of these bins (0.02-0.08Hz, p= 0.03*10^−4^; 0.04-0.16Hz, p= 0.03*10^−4^; 0.08-0.32Hz, p= 0.17*10^−4^; 0.16-0.64Hz, p= 0.12*10^−4^; 0.32-1.28Hz, p= 0.75*10^−4^; 0.64-2.56Hz, p= 0.895*10^−4^; 2.56-4.0Hz, p= 0.035).

**Figure 3.**
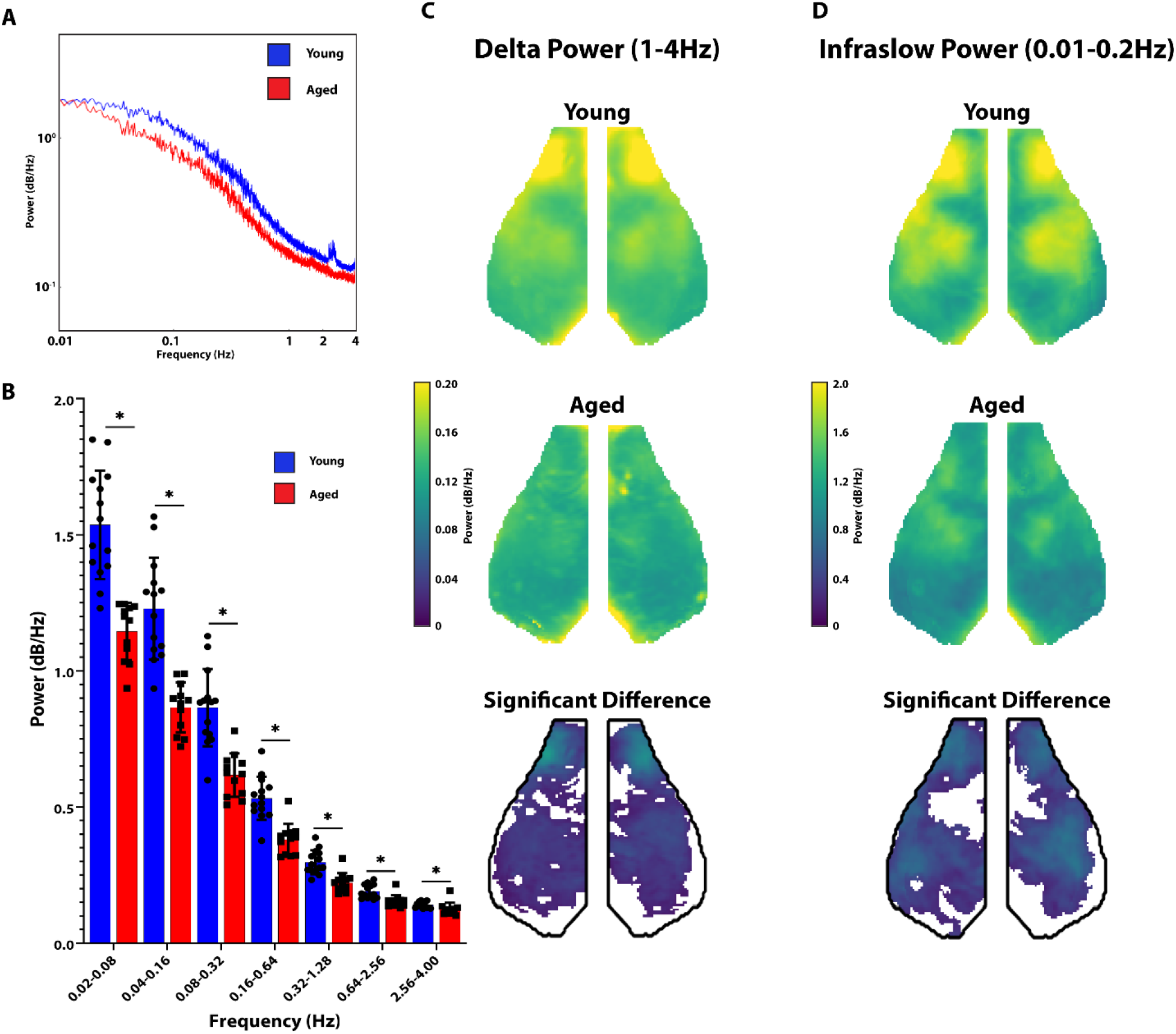
Aged Animals have Significantly Reduced Resting State Power. **A**. Graph of Power as a function of frequency in aged (red) vs young (blue) animals. **B**. Whole brain power divided into octaves in young (blue) vs. aged (red) animals. Young animals had significantly greater whole brain power at all measured frequencies. **C**. Group averaged maps of whole-cortex delta (left) power in young animals (top row) and aged animals (middle row). The bottom row represents the difference between young and aged groups (young-aged) thresholded for significance. **D**. Same as C but in the infraslow band.

We next examined whether the reductions in resting state activity were focal or global. Two canonical frequency ranges were examined, delta (1-4 Hz) and infraslow (0.01-0.2 Hz). Maps of average power in young (top row) and aged (middle row) in both the delta (Figure 3c) and infraslow (Figure 3d) bands are shown. Within both frequency bands, we see the greatest power in the motor, parietal and somatosensory cortices similar to prior observations(36). We then created difference maps between young and aged animals (young minus aged) thresholded for significance. We found a significant reduction in resting state activity in the delta and infraslow frequency bands throughout most of the cortex (Figure 3c and Figure 3d).

### Functional connectivity reductions are network specific in aged mice

Given the broad decline of spontaneous activity, we next examined the effect of aging on network connectivity. 8 canonical “seeds”, representing previously defined network nodes (as described in Methods), were placed on each side of the brain at the mid-point of unique networks (left and right frontal, motor, somatosensory, parietal, cingulate, retrosplenial, auditory, and visual). Group average maps of the connectivity of the entire cortex to the seed networks in the delta (Figure 4a) and infraslow (Figure 4b) frequency bands in young and aged (Figure 4A_1_ and B_1_) animals were generated. Correlation matrices(40, 41) were generated in young and aged mice (Figure 4A_2_ and 4B_2_), and difference matrices were calculated (young minus aged) (bottom row Figure 4A_2_ and 4B_2_). We found that aging was associated with significant deterioration of connectivity between select networks. Most notably, connectivity between parietal and motor and cingulate and motor seeds were significantly different with age across both the infraslow and delta frequency bands. Differences were also observed in retrosplenial and somatosensory seeds. We did not observe connectivity differences within the visual or frontal seeds.

**Figure 4.**
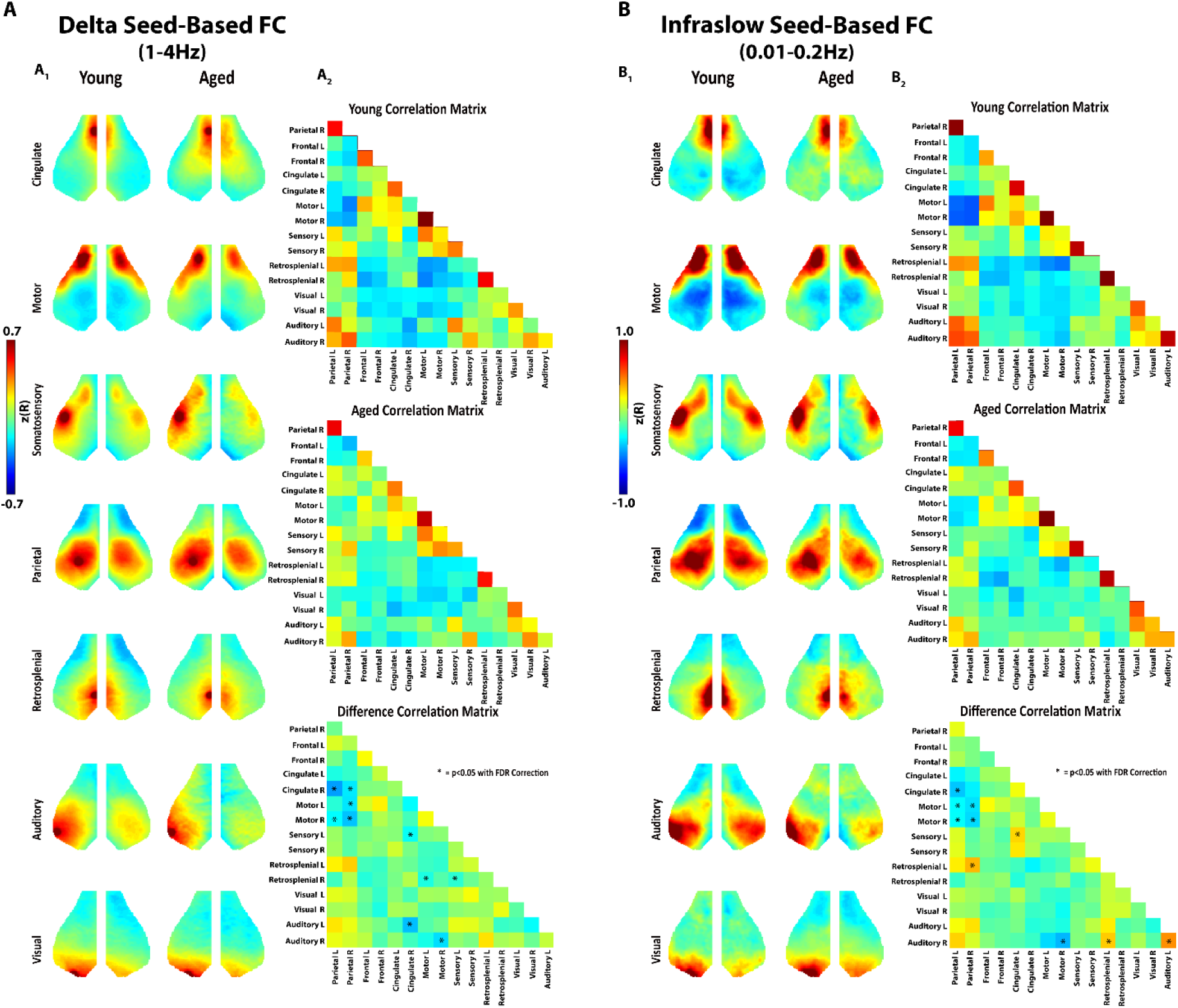
Aged Animals have Significantly Reduced Internetwork Connectivity using Seed Based Analysis. **A**_**1**_. Seed based analysis of functionally connected networks in the mouse cortex in the delta frequency band. Each map represents group average connectivity values for seeds placed in distinct regions (as labeled). The left column is young animals. The right column is older animals. **A**_**2**_. Connectivity correlation matrices for all seeds in young animals (top), aged animals (middle), and the difference between the two (young minus aged) (bottom). Asterisks are placed over cells in the matrix that represent a significant difference (p<0.05 with FDR correction). **B**_**1**_. Seed based analysis of functionally connected networks in the mouse cortex in the infraslow frequency band. Group average maps are shown similar to A. **B**_**2**_. Correlation matrices in the infraslow band similar to A.

Because a large proportion of the FC changes we observed were interhemispheric, we next compared homotopic FC in aged and young mice by measuring the correlation of each pixel on one side of the brain to its contralateral homologue. As this FC measure is symmetric about midline, homotopic FC is visualize in the left hemisphere while network assignments (based on the Paxinos atlas) are visualized in the right hemisphere. Consistent with prior work(41), stronger homotopic FC was observed in motor, somatosensory, and visual cortices in both the delta frequency (Figure 5a) and infraslow frequency bands (Figure 5b). Age-related homotopic FC changes were strikingly focal in comparison to the global power changes. Aged mice showed significantly less homotopic functional connectivity in the motor (delta and infraslow bands) and somatosensory (infraslow band) and visual (infraslow band) cortices.

**Figure 5.**
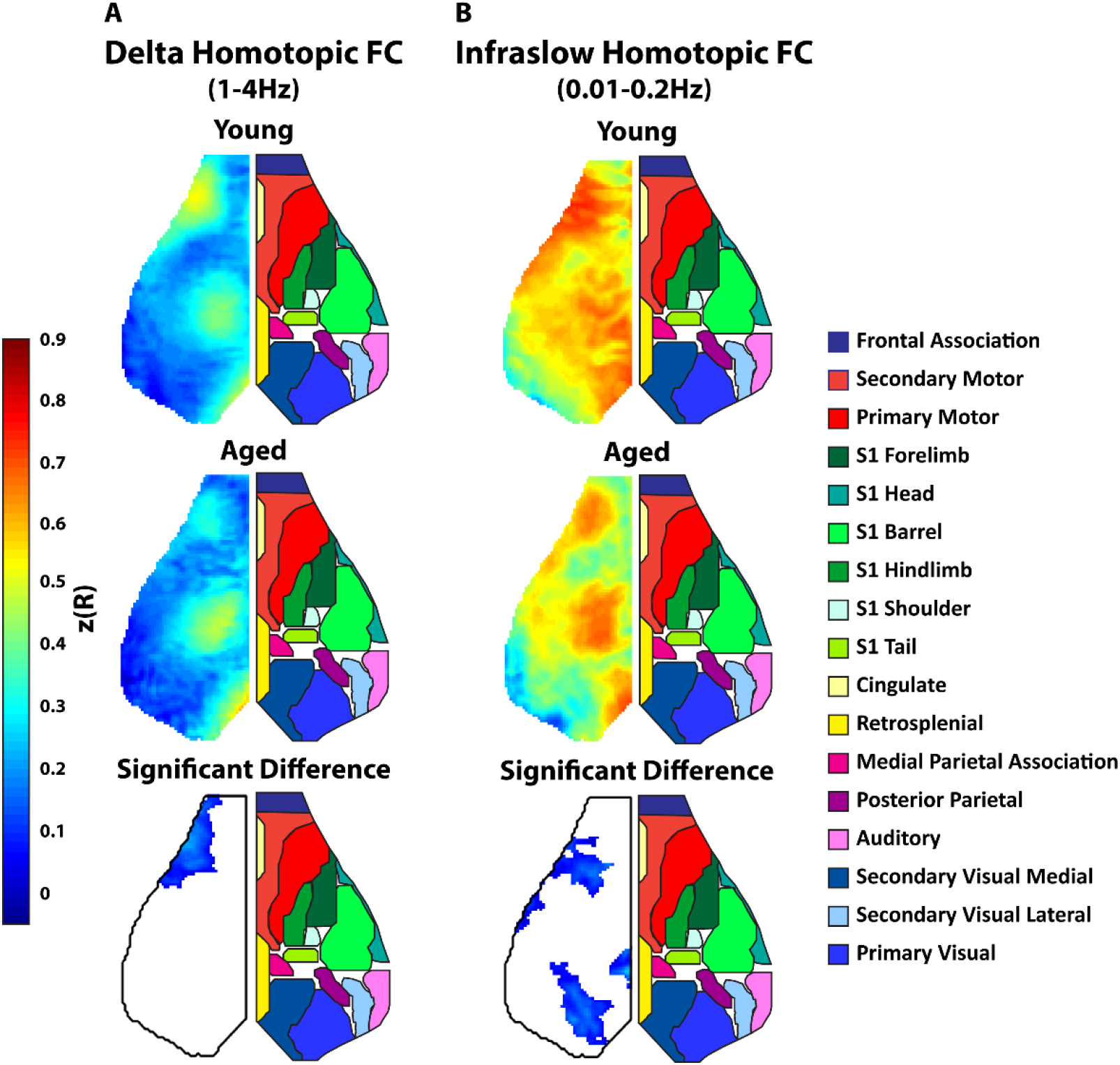
Aged Animals have Significantly Reduced Homotopic Functional Connectivity. **A**. Group averaged maps of whole cortex homotopic FC (correlation between each pixel and its mirror pixel) in the delta (1-4Hz) frequency band. Top map is the average homotopic FC in young animals. The middle map is average homotopic FC in young animals. The bottom map is difference between the average young and aged maps (young-aged) thresholded for significance. Since the maps were symmetric, the left side of the cortex is shown and the right side shows a map of approximate cortical networks based on the Paxinos atlas. **B**. Similar group average maps as **A** but in the infraslow band.

Another salient observation in the seed-based FC maps of Figure 3 was a global trend towards reduced positive correlation in aged mice. We quantified this decline in FC by calculating the total number of functional connections (node degree) of each pixel across the entire cortex in the delta (Figure 6a) and infraslow (Figure 6b) frequency bands. We found robust cortical node degree in both frequency bands in the motor, somatosensory, and visual cortices. We found more wide-spread age-related loss of node degree in the infraslow frequencies than the delta frequency band. Within the delta frequency band, aging was associated with significant reduction of motor cortex node degree. Within the Infraslow band, aging was associated with significant reduction of somatosensory, and visual cortex node degree. In addition to pixel by pixel analysis of node degree across the entire brain (Figure 6) we also examined average node degree within the seeds used for seed-based FC analysis and compared them across young and aged animals (Figure 7). This again demonstrated widespread loss of node degree with aging (more predominant in the infraslow band) particularly in motor, somatosensory, and visual cortices.

**Figure 6.**
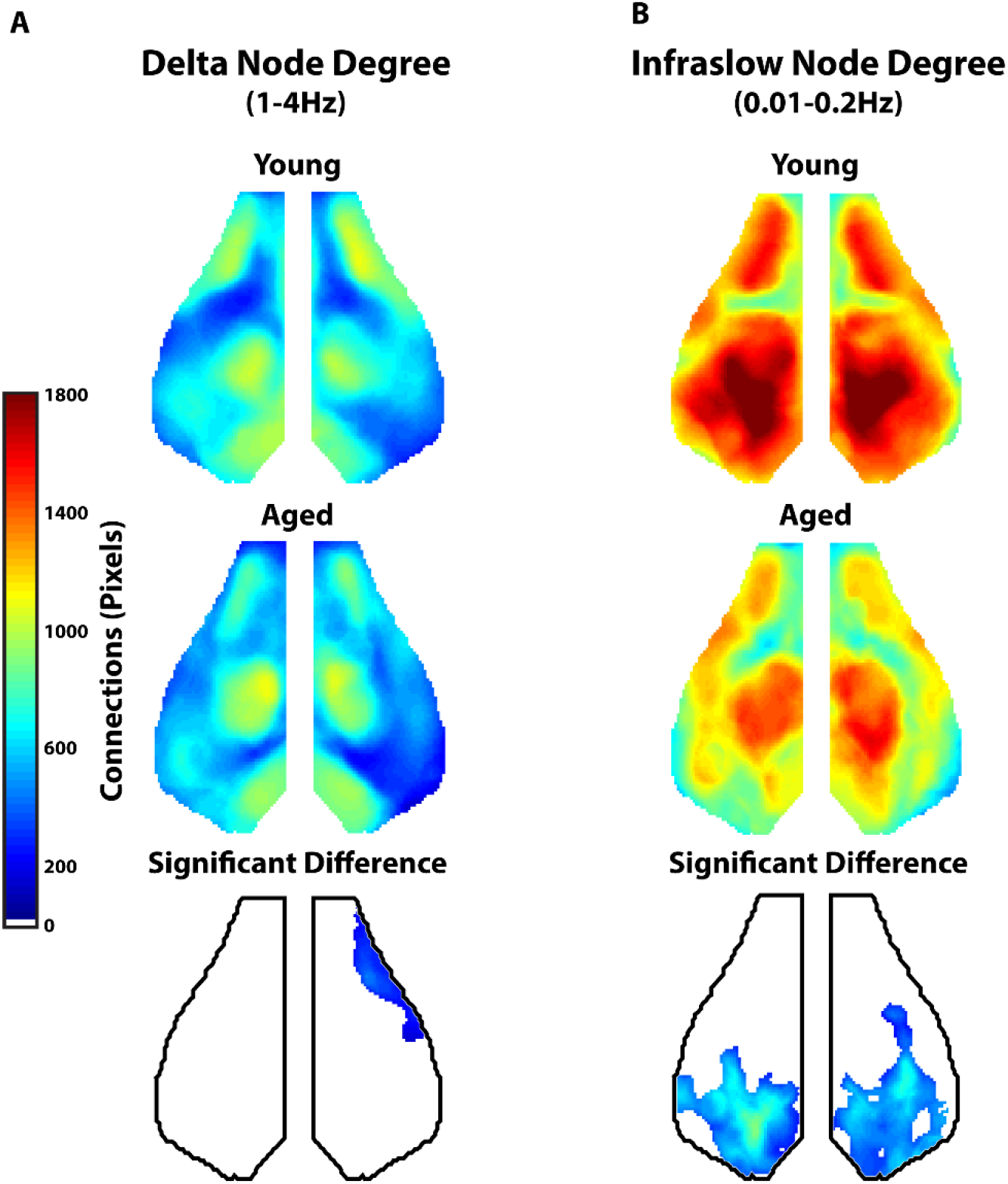
Aged Animals Have Significant Reduction in Global Node Degree. **A**. Group average map of global (whole cortex) node degree in the delta frequency range (1-4Hz) for young (top) and aged (middle) animals. The bottom map represents the difference between young and aged animals in global node degree thresholded for significance. **B**. Group average map of global (whole cortex) node degree in the infraslow frequency range (0.01-0.2Hz) for young (top) and aged (middle) animals. The bottom map represents the difference between young and aged animals in global node degree thresholded for significance.

**Figure 7.**
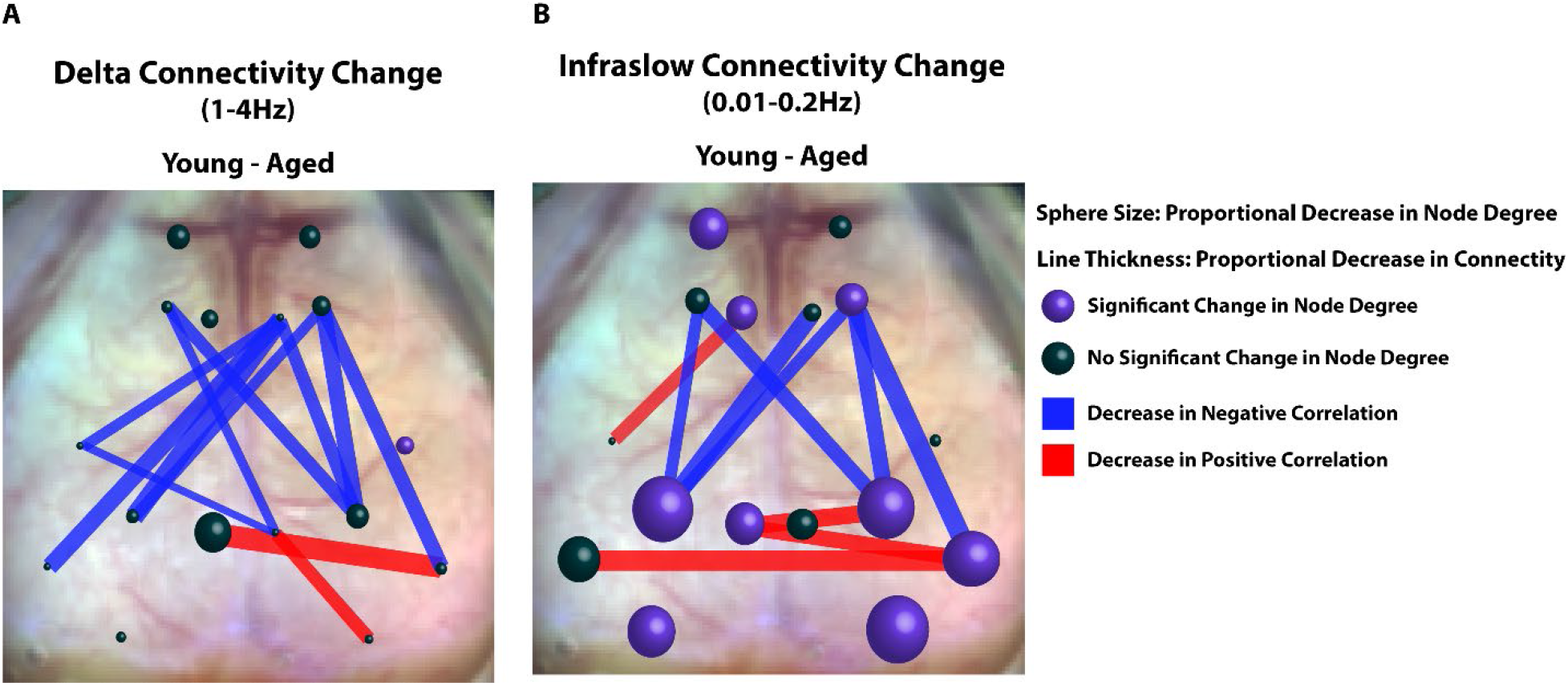
Summary of Connectivity Changes in Aged vs. Young Animals. **A**. Representative diagram of connectivity difference between young and aged animals (young minus aged) within the delta band. Spheres are at locations of seeds used in seed-based FC analysis. Sphere size correlates to the average change in node degree within the seed between young and aged mice (Young - Aged). Average node degree within the seeds was compared between young and aged animals using a student’s T test. Purple spheres represent significant differences in average node degree at that location (p<0.05). Lines between seeds represent significant differences in connectivity (z(R)) between young and aged groups. The thickness of the lines is proportional to magnitude of the correlation difference. Red lines represent age associated decreases in positive correlations and blue lines represent age-associated decreases in negative correlations. **B** Same as A, but in the infraslow band.

In contrast to our observation of a relatively global decrease in resting state power within the aged cohort, we observed more network specific changes in functional connectivity with aging (Summarized in Figure 7). Age-associated deterioration of node degree (sphere size) was more predominant in the infraslow band and demonstrated significant reductions in the bilateral visual and parietal cortices as well as smaller reductions in unilateral frontal, cingulate, retrosplenial, and auditory cortices. Seed-based deterioration of FC (red and blue lines) in aged animals as noted in figure 4 was observed in several networks including sensory, motor, and cingulate networks. Notably, the majority of aged-associated FC deterioration was between negatively correlated regions (blue vs red lines).

### Global power and node degree correlate with activity in young but not aged mice

To compare spontaneous mouse behavior between young and aged groups, we scored time spent on: wall exploration (Figure 8a_1_), self-grooming (Figure 8a_2_), and inactivity (Figure 8a_3_). We observed significantly less wall exploration time in aged mice (% frames exploring) (Young= 4.93 ± 0.53; Aged=2.57 ± 0.43; p=0.0014) and significantly more inactive time (% frames inactive) (Young= 12.91 ± 2.36; Aged=24.83 ± 2.80; p=0.0027). We did not find a significant difference in grooming behavior between young and aged mice (% frames grooming) (Young= 7.94 ± 1.30; Aged=5.20 ± 0.63; p=0.071).

**Figure 8.**
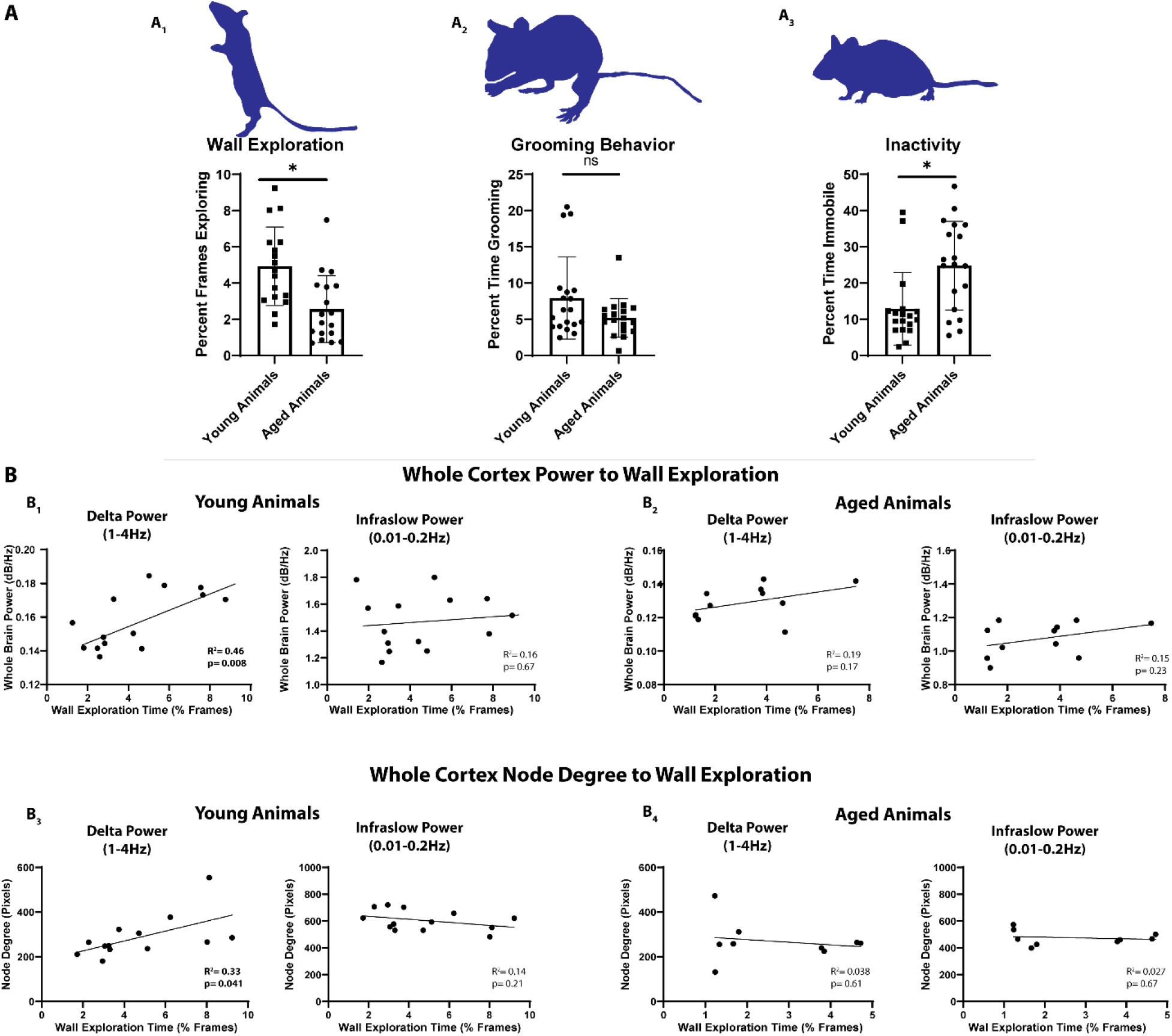
Exploratory Behavior in Young Animals Correlates with Cortical Delta Power and Delta Node Degree. **A**_**1**_. Young animals explored a clear glass container significantly more than aged animals. **A**_**2**_. Grooming behavior was similar in aged and young animals. **A**_**3**_. Aged animals spent significantly more time immobile than young animals. **B**_**1**_. Whole brain power significantly correlated with the degree of wall exploration observed in young animals within the delta but not infraslow frequency band. **B**_**2**_. Whole brain power in aged animals did not correlate with exploration time in either the delta or infraslow frequency band. **B**_**3**_. Whole brain node degree correlated significantly (R^2^=0.43) with the degree of wall exploration within the delta but not infraslow frequency band in young animals. Infraslow node degree did not correlate. **B**_**4**_. There was no correlation between delta or infraslow node degree and exploration in aged animals.

We examined the correlation between whole cortex power in the delta and infraslow spectrum and wall exploration on an individual mouse level. We found that young mice exhibited a significant correlation between wall exploration and whole-cortex, resting-state delta power (Figure 8b_1_) (R^2^ = 0.46; p=0.008). Wall exploration did not correlate with infralow power (Figure 8b_1_) (R^2^ = 0.0.016; p=0.67). We further did not observe a correlation between inactivity and power. Aged mice, which were significantly more immobile overall, did not exhibit this correlation (Figure 8b_2_) (R^2^ = 0.19; p=0.17). We next examined whether whole cortex node degree correlated to mouse activity. We found that average whole-cortex node degree correlated (R^2^ = 0.33; p=0.046) with wall exploration in young mice (Figure 8b_3_), but not aged mice (Figure 8b_4_). As with resting state power, this correlation was noted only in the delta band.

To determine if specific networks were responsible for the correlation between whole brain power/node degree and exploratory behavior, we examined the correlation between exploratory behavior and average resting state power and average node degree within different networks (as outlined in figure 5). Delta power within the somatomotor (R^2^ = 0.3; p=0.042), visual (R^2^ = 0.33; p=0.032), and retrosplenial networks (R^2^ = 0.43; p=0.012) correlated significantly with exploratory activity in young but not aged animals (Supplemental Figure 1). Moreover, Delta band node degree within the somatomotor (R^2^ = 0.32; p=0.04), and visual cortex (R^2^ = 0.33; p=0.04), correlated significantly with behavior in young but not aged animals. Infraslow band power and node degree did not correlate with behavior (Supplemental Figure 2).

## Discussion

We utilized mesoscale imaging in *Thy1*-GCaMP6f mice to directly compare spontaneous neural activity and network connectivity in aged and young animals with high spatiotemporal resolution. Aging was associated with a nearly global decrease in spontaneous network activity across the spectrum of frequencies we examined (0.01-4 Hz). This decrease in resting state power was present throughout most of the cortex. In contrast to the widespread changes in resting state neuronal activity, we observed that aging was associated with more focal changes in functional connectivity. In examining homotopic FC, seed-based FC, and global node degree, aged animals exhibited reduced FC in several cortical regions including motor, parietal, and visual cortices. Examination of spontaneous behaviors revealed that exploratory activity in young mice significantly correlated with whole cortex resting state delta power, but not infraslow power. Aged mice, which were significantly less active, no longer exhibited this correlation. Finally, we observed that whole cortex node degree correlated with exploratory activity in young animals. As with power, this was not observed in the infralsow frequency band and aged mice no longer demonstrated this correlation. Together these data suggest that aging is associated with broad declines in global power as well as local deterioration of connectivity within select cortical networks. These declines correlated with behavioral changes seen in aged mice and may suggest a link between the neuronal and synaptic changes observed in older individuals and behavioral declines associated with aging.

The synchronous activity of networks has been described as a “middle ground” between single neuron activity and behavior(42). Spontaneous neuronal activity across a variety of frequencies has been implicated in a wide range of functions including information transfer, plasticity, memory, attention, and consciousness(19, 20, 43). Importantly, phase synchronization of activity in distinct brain regions is thought to underlie functional connectivity critical for the integration of information(44). Our data show two primary findings with aging: 1) a nearly global decline in resting state activity across a wide frequency range (0.01-4Hz) that extends over much of the cortex (encompassing most of somatosensory, motor, and visual cortices), and 2) more nuanced decline in regional functional connectivity within and across resting state networks (as illustrated in figure 7).

Prior work has shown that aging is associated with significant changes to cortical spontaneous activity. Most relevant is recent magnetoencephalographic work demonstrating widespread linear declines in slow wave EEG power (0.5 -6.5Hz) with aging (22). Compellingly, this work also demonstrated that enhanced delta and theta power was associated with better performance on tests of executive function. Our work adds to the finding that aging is associated a broad decline in spontaneous activity across a wide frequency range including infraslow and delta (0.01-4Hz) and that this decline covers most of the cortex. Additionally, we found that whole cortex delta activity positively correlated with exploratory activity in young animals, but in aged animals (which explored significantly less and had significantly less infraslow and delta power overall), this correlation was no longer observed. Correlations between delta band power and node degree and exploratory behavior suggests a frequency specific role in some spontaneous behaviors which is lost in aged animals who exhibit significantly less exploratory behavior overall.

Observations of age-associated white matter deterioration led to the “disconnection” hypothesis that cognitive decline during normal aging results from changes in the integration of cerebral networks(45). A number of functional imaging studies have since shown that aging is associated with a wide range of network and FC changes(46, 47). Seminal work by Andrews-Hanna et al. demonstrated that aged individuals, free from Alzheimer’s disease pathology, had significantly reduced anterior to posterior functional connectivity within the default mode network(48). Others have further demonstrated that aged individuals have reduced frontoparietal control and default mode network connectivity(49), reduced connectivity within the frontoparietal network(50), and reduced cortico-striatal network FC(51). A recent study using fMRI in mice found an inverse U shape of functional connectivity strength in the sensory motor and default mode networks across the lifespan of mice with connectivity increasing until middle age then declining significantly(52). Here we leverage the high spatial resolution and broad temporal range of mesoscale calcium imaging to show that neural activity in aged mice is globally reduced. We also show that aging in mice is associated with degradation of functional connectivity in discrete cortical networks particularly somatomotor and visual networks. Finally-we show that band specific correlations in power and connectivity with behavior in young, but not aged mice.

Many of the reductions in FC we observed in our seed-based analysis were loss of anti-correlation (summarized by the blue lines in Figure 7a and Figure 7b). This resulted in less discrete network borders (as seen in figure 4A_1_ and 4B_1_). Decreased functional specialization and network integrity has been well described with aging. Hippocampal place cells firing become less specific as animals age(53). Older individuals utilize a greater cortical area to process visual information(54), utilize significantly less cortical specialization when viewing distinct categories of visual stimuli(55), and recruit additional cerebral area during verbal memory encoding(56). Additionally, resting state FC analysis has shown decreased segregation of functional brain systems in aged individuals-particularly those older than 50(57). The diminished anticorrelation and degradation of distinct network borders observed in our study may represent network desegregation with aging in mice. Anticorrelation between networks is thought to be an important component of attention demanding tasks(58) and, of note, human studies have also demonstrated that anticorrelation between the dorsal attention and default mode networks is attenuated in older adults(59).

Recent work demonstrating that delta and infraslow activity likely represent unique neural processes with separate propagation patterns(60) supports our finding. If activity within these frequencies represent unique processing-then it is interesting to consider that while aging appears to universally drive down cortical network activity, its effect on delta activity may disproportionately affect certain somatomotor functions. Delta activity predominates cortical function during development(61) and slow wave sleep(62).

Relevant to our finding that delta activity correlates with exploratory activity, it has been highly implicated in motivational behavior(63, 64). Declines in delta activity with aging may therefore underlie diminished drive to explore a new environment. Delta activity has also been associated with visual attention(65), hunger(66, 67), and reward craving(68), processes to which exploratory behavior would seem closely related. Further delineating possible frequency specific correlations with other behaviors will be an important subject of future work.

Our observation that delta frequency power and node degree correlate to exploratory behavior in young but not aged mice (which exhibit significantly less delta power and exploratory behavior overall) adds to the hypothesis that network dysfunction contributes to age related functional decline. Prior human studies have shown a strong correlation between network behavior and behavioral declines. The majority of this work has focused on age-associated cognitive decline. Default mode network functional connectivity in older adults free of Alzheimer’s pathology has a linear correlation with composite scores of executive function, memory recall, and processing speed(48). Memory decline itself is conversely correlated with default mode network FC(69). Decreased FC within the salience network is noted with aging and the degree of impairment correlates with decreased scores on a battery of cognitive neuropsychiatric tests(70). Innovative work using machine learning has shown that the connectivity profile between the salience and visual or anterior default mode can be used to predict age(71). Healthy older individuals have reduced fronto-parietal and fronto-occiptial connectivity and lower regional clustering scores were correlated with diminished verbal and visual memory scores(72). A study of working memory demonstrated healthy aged individuals had reduced default mode connectivity and that the degree of connectivity between the posterior cingulate cortex and medial prefrontal cortex correlated with memory performance(73). Human imaging studies have also noted a variety age-associated changes in network activation related to motor activity(74) and learning(75). Our study found the greatest differences in connectivity and power to be in the somatomotor networks, therefore we focused our work to this region.

Group-level differences in neural activity and functional connectivity between young and old mice are likely a result of changes in patterns of intrinsic functional organization that naturally occur during normal aging, rather than as a result of changing GCaMP expression. Importantly, we did not find a significant difference in GCaMP positive cell count in aged animals, nor did we detect any of the characteristic signs of cellular stress or toxicity in this or our previous work(76) (e.g. beading of axons, punctate expression of aggregated GCaMP, nuclear expression, or cellular atrophy). Additionally, stimulus-evoked responses were similar across aged and young animals, suggesting that the brain’s ability to process sensory input was comparable across groups. Global reductions in signal power could result in reduced signal coherence, manifesting as lower values of zero-lag correlation. However, changes in FC were region specific while the very broad reductions in band-limited power involved the majority of the cortex. FC differences localized to select networks suggests that our measurements are sufficiently sensitive to detecting age-related alterations in functional brain organization processes.

In conclusion, the data presented here suggest that aging is associated with global declines in resting state activity ranging over a wide range of frequencies and over large portions of the brain. In contrast, aging is associated with focal deterioration of cortical network connectivity most prominent in motor and somatosensory cortex. Finally, we report that whole cortex delta band activity and node degree connectivity correlated with exploratory activity in young, but not aged mice. These data suggest that somatomotor network degradation driving diminished delta activity may disproportionally influence behavioral deficits with aging.

## Methods

### Animals

A total of 33 transgenic *Thy1*-GCaMP6f mice (Jackson Laboratories C57BL/6J-Tg (Thy1-GCaMP6f)GP5.5Dkim/J; Stock Number 024276) were used in this study. 16 (7 female and 9 male) young mice aged (2-3 months) and 17 (8 female and 9 male) aged mice (17-18 months) were included in the study. Mice were housed in standard cages with *ad libitum* access to food and water in a dedicated animal facility under a 12 hour light/12 hour dark cycle. All experimental protocols were approved by the Animal Studies Committee at Washington University and all studies were conducted in accordance with the US Public Health Service’s Policy on Humane Care and Use of laboratory animals.

### Extracranial Window Placement

Mice were anesthetized with inhalation isoflurane (4.0% induction, 1.5% maintenance) and placed into a stereotaxic head fixation frame (Kopf Instruments, Tujunga, CA, USA). Each mouse was placed on a heating pad and temperature was maintained at 37°C via feedback from a rectal probe (mTCII Cell Microcontrols, Norfolk, VA, USA). The head was thoroughly shaved and cleaned and a midline incision was performed as previously described(36, 37, 76). The scalp was retracted, and a custom-made clear Plexiglas window was affixed directly to the skull using dental cement (C & B Metabond, Parkell, Edgewood, NY, USA). Mice were monitored daily after window placement.

### Imaging Acquisition

GCaMP calcium imaging was performed as previously described^29^. Mice were placed in a felt pouch and their heads secured via small screws in the plexiglass windows under an overhead cooled, frame-transfer EMCCD camera (iXon 897, Andor Technologies, Belfast, Northern Ireland, United Kingdom) with an approximately 1cm^2^ field of view covering the entire dorsal view of the brain. Sequential illumination was provided by a 454 nm blue LED (Mightex Systems, Pleasanton California, USA). An 85 mm *f*/1.4 camera lens was utilized (Rokinon, New York, New York, USA) and images were acquired at a frame rate of 16.81 Hz. The CCD was binned at 4 × 4 pixels to increase SNR giving a final resolution of 128 × 128 pixels. The LED and the CCD were synchronized and triggered via a DAQ (PCI-6733, National Instruments, Austin, TX, USA) using custom written MATLAB scripts (MathWorks, Natick, MA, USA). Data from all imaging sessions were acquired and stored in 5-minute intervals.

For awake, resting state imaging, mice were acclimatized to the imaging felt pouch for several minutes and allowed to move freely with the exception of their restrained head. The room was kept dark and imaging was only commenced when mice were resting comfortably. For stimulus induced imaging, mice were anesthetized with an intraperitoneal injection of ketamine/xylazine (86.9mg/kg ketamine, 13.4mg/kg xylazine). After anesthesia, the animals were placed on heating pads maintained at 37°C as described above. Transcutaneous forepaw stimulation was performed similar to prior studies(38) via placement of microvascular clips (Roboz Srugical Equipment, Gaithersburg, MD, USA) on the right wrist. Electrical stimulation was via a block design (0.5mA, 3hz stimulation, 5 seconds, 30 events in 5 minutes).

### Image Processing

All data was subjected to an initial quality check and all data with movement contamination were excluded. Image processing was as previously described(37, 38, 40). A binary brain mask was created and applied by tracing the field of view to include only the brain component using the roipoly.m function in Matlab. Image sequences from each mouse were affine-transformed to a common atlas space (based on the Paxinos mouse atlas) using the positions of bregma and lambda. Images were smoothed with a 5 × 5 Gaussian filter. Reflectance Changes in the 454nm reflectance were used to analyze the GCaMP signal and a ratiometric correction for absorption by hemoglobin and deoxyhemoglobin was used as previously described^32^. The time traces for all pixels within the brain mask were averaged to create a global brain signal. For all resting state and activation analysis, this global signal was regressed from every pixel’s time trace to remove global sources of variance. Resting state data was filtered into canonical frequency ranges of 0.01-0.2 Hz (Infraslow) and 1-4Hz (Delta). Power spectral analysis was computed using a Hann window and fast fourier transform. The square moduli of these FFTs were then averaged across the brain to produce the final power spectra.

Seed-based functional connectivity was calculated by placing seeds (0.25mm diameter containing approximately 30 pixels) in each hemisphere (16 total seeds). Seeds were placed at canonical locations expected to correspond with the bilateral frontal, motor, somatosensory, parietal, cingulate, retrosplenial, auditory, and visual cortices. Seed locations were determined prior to imaging using an anatomic atlas(77) similar to our prior work(40, 78, 79). For each seed region, time traces of GCaMP fluorescence were correlated with time courses in all brain pixels generating an FC map for each seed. Individual mouse maps were transformed into Fisher z-scores and averaged within each group (aged and young). Homotopic functional connectivity was calculated by correlating the time trace of each pixel in the left hemisphere to the time trace of its contralateral homolog. Correlation values underwent Fisher z-transformation and were averaged on a group level. Node degree was calculated by determining the number of pixels (degree) with which a particular pixel (node) had a correlation above threshold (r ≥0.4) similar to prior work(41).

### Histology

Mice were deeply anesthetized with FatalPlus (Vortech Pharmaceuticals, Dearborn, MI, USA) and transcardially perfused with 0.01 mol/L, ice cold, heparinized, Phosphate Buffered Saline (PBS). The brains were removed and submerged in 4% paraformaldehyde for 24 hours. Brains were then transferred to a solution of 30% sucrose in PBS Brains were sliced at 50µM thickness on a sliding microtome (Microm, Boise, ID, USA). Sections were stored in 0.2M PBS, 20% Sucrose, and 30% ethylene glycol at -20°C. Imaging of GCaMP6f expression was performed using an inverted confocal microscope (Nikon A1-Rsi). Images were exported and fluorescent (GCaMP+) cells were counted manually (ImageJ).

### Evaluating Spontaneous Behavior

Spontaneous mouse activity was observed by placing the mice in a clean, clear container and recording their spontaneous behavior for 5 minutes. Cameras (Kodak, Rochester, NY, USA) were placed immediately adjacent to the cylinder with a side view and recorded for 5 minutes (60FPS). Videos were exported and analyzed using custom-written scoring software on a frame-by-frame basis. We divided spontaneous mouse activity into 3 categories for simpler quantification. Wall exploration was quantified as the number of frames the animals was touching the walls of the container normalized to the total number of frames. Self-grooming behavior was defined as rubbing paws, licking, scratching, and running paws through hair normalized to total number of frames. Immobility was defined as the number of frames not moving normalized to the total number of frames.

### Statistics

All statistical analyses were performed using either Matlab or Prism Graphpad Software. For evoked responses, an ROI was drawn around the area of somatosensory activation (defined by all the pixels with activation amplitudes during forepaw stimulation within the 75% of the maximum activation). Within this ROI, we calculated the average intensity in aged and young animals. Additionally, we calculated total area of activation between aged and young animals by counting the number of pixels within the defined 75^th^ percentile of the maximum activation. Statistically significant differences in whole brain power, node degree, and homotopic functional connectivity were calculated by comparing young and aged mice with unpaired, two sample *t* tests.

Analysis of power and FC carries a high risk of Type 1 error due multiple comparisons across the large number of pixels and the risk of Type 2 error when applying a classic correction such as the Bonferroni across the mask. We used a cluster-based, non-parametric, permutation method described by Maris and Oostenvel(80), which groups pixels into clusters based on data variance across conditions, thus minimizing the number of multiple comparisons and optimizing both Type 1 and Type 2 errors. The test statistics were *t* test statistics for z(R), power, and node degree. Briefly, pixels with test statistics less than p value 0.05 threshold were clustered into continuous areas, from which the cluster statistics were calculated by the summation of the test statistics of the pixels in each cluster. A null hypothesis distribution of cluster statistics was obtained by iterating 2000 times a random assignment of data to different conditions, calculating the test statistics of each pixel, and obtaining the cluster statistics. Clusters were determined to be statistically significant if they were greater than 97.5 percent of the null hypothesis cluster statistic distribution with n degrees of freedom (where n= total subject/2). Difference maps were generated for and thresholded to significance for power, homotopic FC, and node degree.

Seed based functional connectivity matrices were compared via an unpaired Student’s t test with a false discovery rate (FDR) correction. Behavioral correlations were performed by plotting exploration against node degree or power (delta or infraslow) within a region of interest drawn around the cortical space which showed significant aged-related differences and performing simple linear regression to calculate an R squared value. Correlations were tested for non-linearity via an F test and a p value of less than 0.05 was considered significant.

## Acknowledgements

This work was supported by National Institute of Health grants R37NS110699 (JML), R01NS084028 (JML), R01NS094692 (JML), RO1-NS102870 (AQB), K25-NS083754 (AQB), R01NS078223 (JPC), P01NS080675 (JPC), F31NS103275 (ZPR), K08-NS109292-01A1 (ECL), as well as the McDonnell Center for Systems Neuroscience (AQB). This work was also supported by American Heart Association Grants 20CDA35310845 (AJA) and 20CDA35310607 (ECL). We thank Karen Smith for her help with behavior acquisition and animal care.

---

**Supplemental Figure 1.**
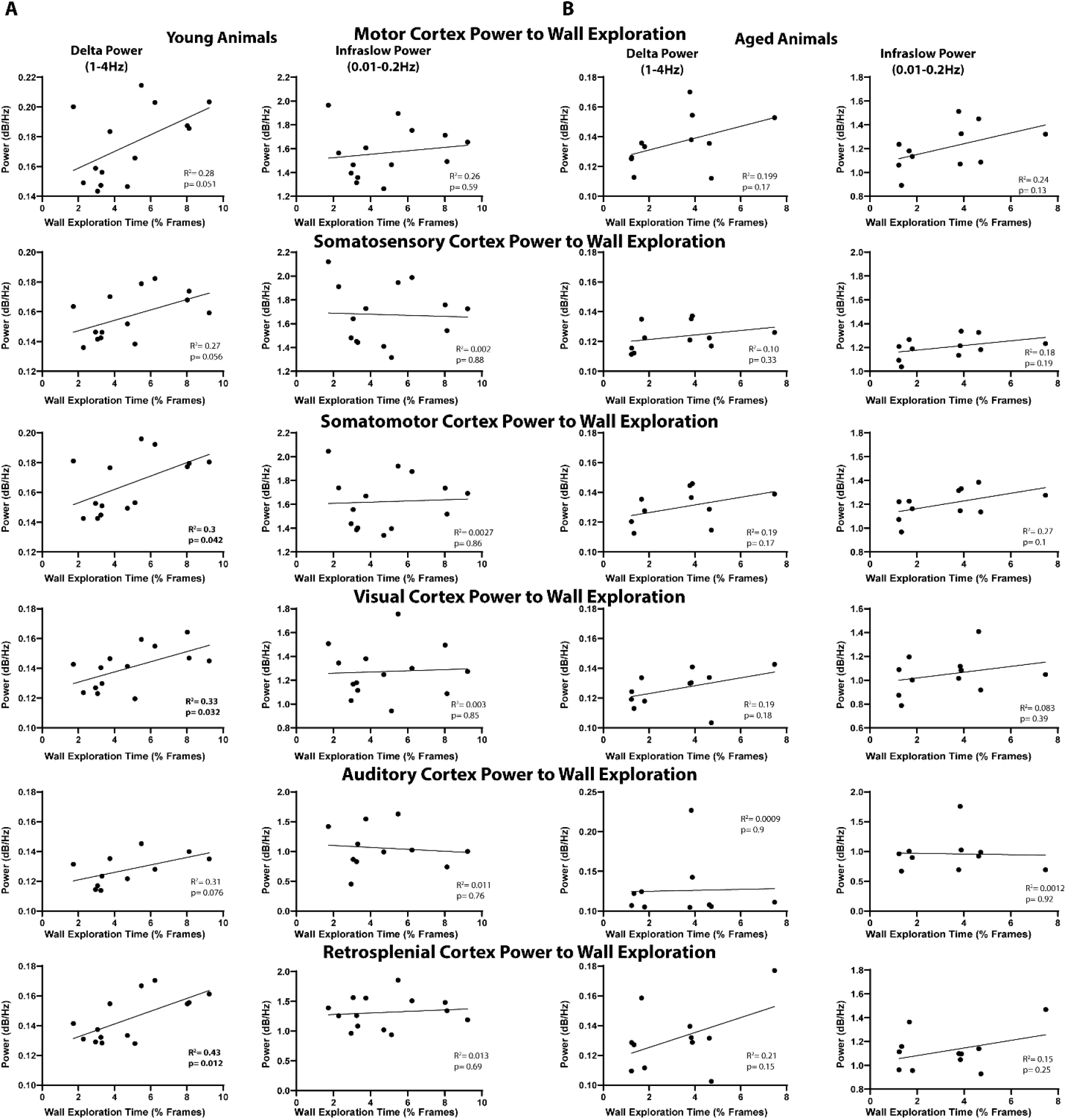
Correlation Between Exploratory Behavior and Cortical Power in Select Network. **A**. Summary of correlations between exploratory behavior and power in the motor cortex (first row), somatosensory cortex (second row), combined somatomotor cortex (third row), visual cortex (fourth row), auditory cortex (fifth row), and retrosplenial cortex (sixth row) in young animals. The first column represents delta power and the second column represents infraslow power. Somatomotor (R^2^=0.3), visual (R^2^=0.33), and retrosplenial (R^2^=0.43) cortices correlated significantly with exploratory behavior in the delta band. There were no correlations in the infraslow frequency range. **B**. Summary of correlations between exploratory behavior and power in aged mice. No significant correlations were observed in the aged group.

**Supplemental Figure 2.**
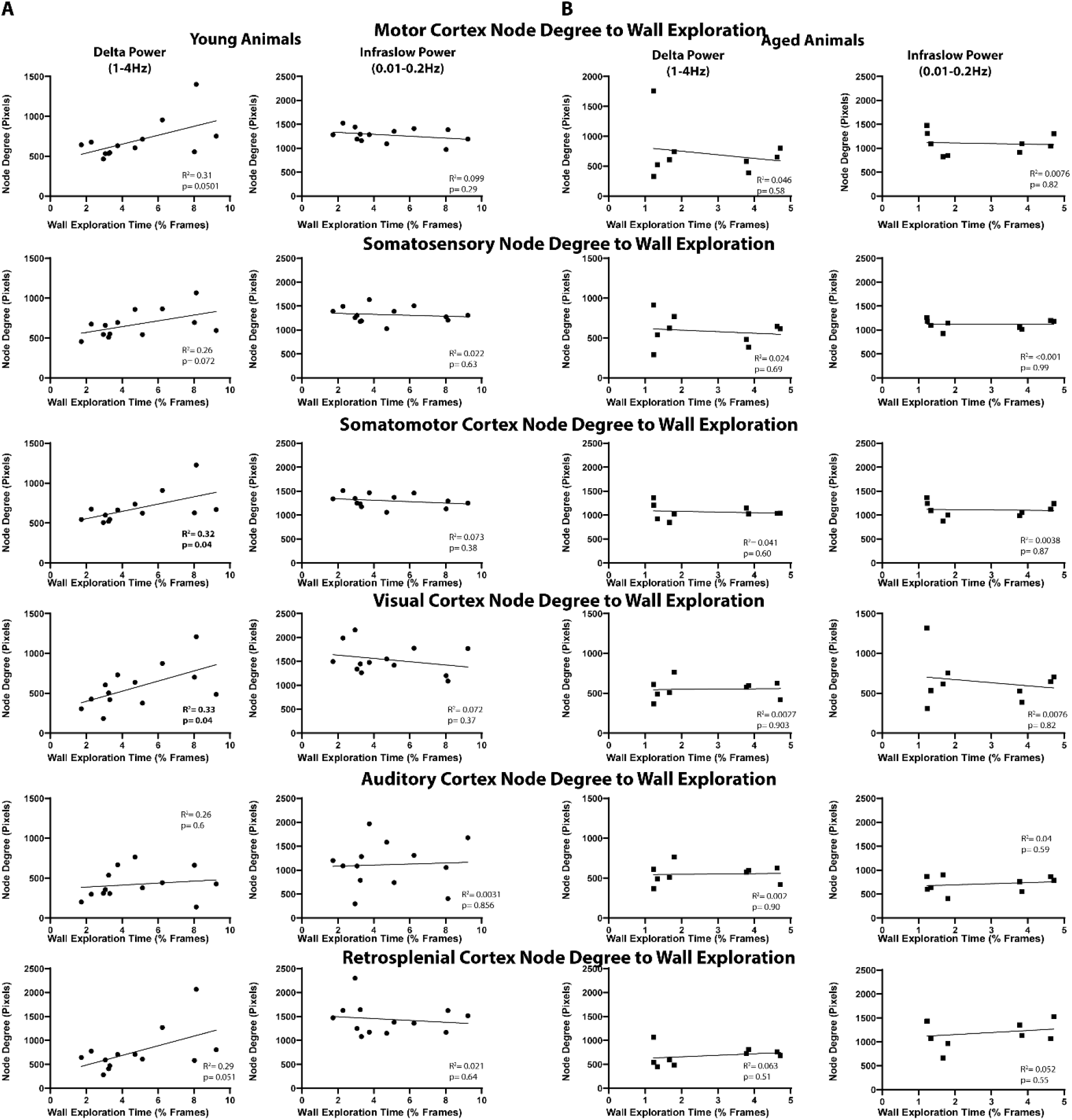
Correlation Between Exploratory Behavior and Cortical Power in Select Network. **A**. Summary of correlations between exploratory behavior and average node degree in the motor cortex (first row), somatosensory cortex (second row), combined somatomotor cortex (third row), visual cortex (fourth row), auditory cortex (fifth row), and retrosplenial cortex (sixth row) in young animals. The first column represents node degree in the delta band and the second column represents node degree in the infraslow band. Somatomotor (R^2^=0.32), visual (R^2^=0.33) cortices correlated significantly with exploratory behavior in the delta band. There were no correlations in the infraslow frequency range. **B**. Summary of correlations between exploratory behavior and node degree in aged mice. No significant correlations were observed in the aged group.

